# Comparison of the bacterial and methanotrophic diversities between an Italian paddy field and its neighboring meadow

**DOI:** 10.1101/535229

**Authors:** Mohammad Ghashghavi, Eric R. Hester, Viktoria Oliver, Claudia Lüke, Mike S. M. Jetten, Sebastian Lücker

## Abstract

Methane is a potent greenhouse gas that contributes to global warming. However, under certain conditions, its release into the atmosphere can be mitigated by methane-oxidizing microorganisms. Typically, cultivated wetlands (i.e., paddy fields) are a major source of methane (CH_4_) while forests and meadow uplands are considered to be CH_4_ sinks. As the global need for rice production increases each year, more uplands are converted to inundated paddy fields. To investigate soils that may be converted into productive land for rice production, we investigated a paddy field and adjacent meadow in Northern Italy. Using a combination of 16S rRNA gene amplicon sequencing to analyze the bacterial community, and gas flux measurements to quantify CH_4_ emissions, we looked for differences between classically defined CH_4_ sinks (meadow soils) and CH_4_ sources (paddy fields). Analysis of the total bacterial community revealed that the family Fimbriimonadaceae, which belongs to the phylum Armatimonadetes, was significantly higher in paddy field soils driving the difference between paddy and meadow soils. Whereas, we found that the methylotrophic families Methyloligellaceae and Methylomirabilaceae were also present in higher relative abundance in the paddy field. Despite these major differences, CH_4_ fluxes were highly variable between the two sites with no significant differences observed. Furthermore, we found the Methylomonaceae family to be more abundant at the center of a neighboring paddy field compared to the edge of the paddy field from the current study, hinting at methanotrophic variation based on location. Taking these results into account, we propose a conceptual model to explain possible scenarios that may result in paddy and meadow fields not exhibiting classical source/sink properties. These findings call for caution when including paddy and meadow areas separately into global CH_4_ flux calculations, and urge further research to discern drivers of CH_4_ cycling under a range of environmental conditions rather than relying on assumptions.

## Introduction

One of the most abundant greenhouse gases in the Earth’s atmosphere is methane (CH_4_), with its concentration steadily increasing as a result of anthropogenic activities (Wahlen, 1993; Dean *et al*., 2018). Approximately 40% of the sources of atmospheric CH_4_ are represented by natural and cultivated wetlands. In order to meet the global need for food, additional land is being converted into cultivated wetlands including inundated paddy soils. This has resulted in an increase in their total area by approximately 1% annually (IPCC, 2013; Pearman, 1986; Dlugokencky *et al*., 1994). Heavily fertilized fields for rice production are considered to be among the highest sources of CH_4_ emission on the planet (Lowe, 2006). The expansion of rice production in combination with increasing global temperatures could exacerbate CH_4_ emissions from these source environments (Aselmann & Crutzen, 1989; Conrad, 2009). Understanding the potential factors that control CH_4_ emissions from upland and wetland environments is necessary to accurately predict future atmospheric concentrations of CH_4_.

In wetlands, CH_4_ emission to the atmosphere results from a greater production of CH_4_ by methanogenic microorganisms than what is oxidized by methanotrophic microorganisms (Cao *et al*., 1998). Although some CH_4_ in paddy soils can be oxidized under anoxic conditions at the expense of nitrate by *Methanoperedens* archaea (Vaksmaa *et al*., 2016; Welte *et al*., 2016), most methane seems to be oxidized by aerobic methane-oxidizing bacteria (MOB) (Lüke *et al*., 2010). Known MOB belong to the *Proteobacteria, Verrucomicrobia* and NC10 phyla (Hanson & Hanson, 1996; Wu *et al*., 2011; Op den Camp *et al*., 2009). Within the *Proteobacteria*, methanotrophs are further classified into two main types known as type I and type II (Hanson and Hanson, 1996). Type I methanotrophs are affiliated with the *Gammaproteobacteria* and assimilate carbon via the ribulose monophosphate (RuMP) pathway, while type II belong to the *Alphaproteobacteria* and utilize the serine pathway for carbon fixation. These two types of MOB were shown to be the dominant methanotrophs in paddy fields, with their growth and activity influenced by soil conditions (i.e., organic content, pH, temperature), fertilizer application and vegetation cover (Hanson and Hanson, 1996; Zheng *et al*., 2008).

Although previous studies have focused on the effect of various agricultural practices, climate change and soil features on methanotrophs, most have only included a wetland (i.e. methane source; for references refer to supplementary table S2) or an upland (i.e. methane sink; for references refer to supplementary table S2). Only few included both environments in their experimental design (Skov et al., 2017; Hondula and Palmer, 2017). This has left the literature split between what main factors are influencing methanotrophic community structure within these environments. Therefore, it is important to further investigate differences in the microbial community and CH_4_ fluxes of these two environments to better understand the contributions to methane emissions and global climate change.

In this study, we investigated the bacterial community from a paddy field and an adjacent meadow by 16S rRNA gene amplicon sequencing. In order to identify any influence soil cultivation has induced on the soil bacterial community, we compared the bacterial community, with special interest in the methanotrophic community, paired with CH_4_ fluxes from a neighboring meadow. We observed that the paddy field and the meadow had distinct bacterial communities being driven by the families Fimbriimonadaceae and Methyloligellaceae along with other methanotrophic groups, although the CH_4_ fluxes did not differ significantly between the classical CH_4_ source (paddy field) and sink (meadow) soils and were found to be highly variable across both environments. By combining the findings of this study with previous literature, we propose a conceptual model that provides several explanatory scenarios for these soils not exhibiting behaviors assumed to be universal to paddy and meadow soils.

## Materials and Methods

### Study site

Field experiments took place at the rice research facility Vercelli, Italy (45°19’25.6’N 8°22’14.2”E). This field has been under rice cultivation with the rice variety *Oryza sativa* temperate japonica Onice for the last 30 years, with irrigation waters coming from the river Sesia during the growing season (May – September) and fields left fallow during the winter months. Sample acquisition took place during the maturing stage of the rice plants. The neighboring meadow has been left uncultivated for the last 30 years, contains sandy soil, and is covered with grass and small bushes. Data obtained from the study by Vaksmaa and colleagues (08°22′25.89″E; 45°19′26.98″N) is from a neighboring paddy field that has gone through the same farming practices and planted with the same rice cultivar, *Oryza sativa*.

### Soil-atmosphere gas exchange and environmental variables

Soil-atmosphere CH_4_ exchange was determined using a static chamber approach (Livingston *et al*. 2005) in July 2015 from the rice paddy field and meadow at the Italian Rice Research Unit in Vercelli, Italy. Measurements were made by using a 10 L volume PVC cylindrical flux chamber, covered with a gas tight lid. Chambers were fitted with small computer fans to promote even air mixing (Pumpanen *et al*., 2004) and a small vent to prevent pressure changes inside the chamber as air was extracted (Hutchinson & Livingston, 2001). Temperature was measured by fitting a temperature probe in a small hole made at the top of the chamber. At the time of sampling, six chambers were placed carefully on the ground, where an airtight seal was created due to the permanently standing water. On the meadow site, where there was no water, a gas tight seal was created by fixing a rubber skirt to the bottom of the chambers. The headspace samples were collected from each flux chamber at five intervals over a 35-minute enclosure period using a gas tight syringe and 1 meter of tygon tubing, in order to prevent disturbance while sampling. Gas samples were stored in pre-evacuated Exetainers^®^ (Labco Ltd., Lampeter, UK), shipped to the University of Aberdeen, UK and subsequently analyzed for CH_4_ concentrations using an Agilent 6890 series gas chromatography system, with a single flame ionization detector (FID) for CH_4_. Repeated analysis of standards determined that instrumental precision error was <10%.

Flux rates were determined using the HMR package (Pedersen *et al*., 2010) in R 3.0.2 (R Core Team 2012) by plotting the best-fit lines to the data for headspace concentration (ppm) against time (minutes) for individual flux chambers. The Ideal Gas Law was used to convert gas concentrations (ppm) to molar concentrations. Fluxes were then reported in mg CH_4_-C m^−2^ hr^−1^. Soil temperature (at 10 cm) was simultaneously measured in three locations adjacent to the chambers using a type K thermacouple (Hanna Instruments Ltd., UK).

### Sample acquisition and DNA Isolation

A total of 21 soil samples were collected near the edge of the paddy field that is neighboring a meadow separated by a ditch located in the described study site. As seen in the sampling diagram (Figure S1), a total of 6 sites at the edge of the paddy field were used for samples to be collected using a 10 cm metallic core with a diameter of 8 cm. Samples were taken at three, six and nine meters from the meadow. Each row of samples was also three meters apart. To exclude rhizospheric soil, each sample site was chosen carefully to be equally distant from all neighboring rice plants, and contain no visible root material from the plants. The soil slurry from the cores was then mixed thoroughly and transferred to 50 mL falcon tubes. Lastly, 15 sampling sites within the meadow were used as described in figure S1. The samples were taken at three, six and nine meters away from the paddy field by using a 50mL falcon tube containing the top 7cm of the dry soil. Each row of samples within the meadow was also three meters apart. Soil surface was cleared before sampling to only include bulk soil. All paddy and meadow samples were stored at −20 °C for further analysis. Bulk soil cores obtained by Vaksmaa and colleagues from the center of a neighboring paddy field were done using an 80-cm soil augers at 5-m intervals (Vaksmaa et al., 2017). The cores were later divided up at each 5-cm depth, however, we only compared the data obtained from the top 10-cm.

DNA from all samples was extracted using the MO BIO Power soil isolation kit following the manufacturer’s protocol (MO BIO Laboratories, USA) with one modification. In the mechanical cell lysis step, the soil samples were beaten with glass beads at 30 s^−1^ frequency for 1 minute using a MO BIO 96 well plate shaker. The quality and quantity of the DNA was checked using gel electrophoresis and spectrophotometric analysis (NanoDrop 1000, Thermo Scientific, USA).

### 16S rRNA gene amplicon sequencing

A two-step PCR protocol adapted from Klindworth *et al*., 2012 and Berry *et al*., 2012 was used to amplify bacterial 16S rRNA gene sequences using the universal bacterial primers B341f and B785r. The PCR program started an initial denaturation step at 98°C for 10 minutes, followed by 25 cycles of denaturation at 95°C for 1 minute, annealing at 60°C for 1 minute, extension at 72°C for 2 minutes, followed by a final extension step at 72°C for 10 minutes. The PCR products were purified using QIAquick PCR product purification kit following manufacturer’s protocol (Qiagen Inc., Germany). Per sample, a total of six PCR reactions were performed. All samples were checked for quality and quantity with gel electrophoresis and fluorescence-based analysis (Qubit 2.0, Thermo Fisher Scientific, USA) before being used in a second PCR step with barcoded B341 forward primer and reverse B785 primer containing the P1 adapter. The PCR conditions were as described above, but only 10 cycles were performed. Subsequently, all parallel reactions were pooled, purified and checked for quality again as described above. Each pooled sample was then further analyzed for quality and quantity in the last step using a Bioanalyzer (Agilent Technologies) following the manufacturer’s protocol and sequenced on the Ion Torrent PGM (Thermo Fischer Scientific).

### Bioinformatic analysis

Raw reads from the Ion Torrent run were analyzed using mothur (Schloss *et al*., 2009). The workflow consisted of eight steps: file processing, quality-based trimming, alignment based processing, pre-clustering, chimera removal, contamination removal, OTU clustering and generation of OTU files. The quality-based trimming step was done with parameters set as follows: PDIFF = 2, MAXHOMOP= 8, MAXAMBIG – 0, QWINDOWAVERAGE =20, QWINDOWSIZE = 50, MINLENGTH = 200 and MAXLENGTH = 450. In the alignment based processing step, the Silva database (v132) (Quast et al., 2012) was used for the DB_ALIGN and DB_TAX function with these parameters: OPTIMIZE = “start-end”; CRITERIA = 95. Lastly, pre-clustering parameter (DIFFS = 2), contamination removal parameter (CLASS_CUTOFF = 80), distance matrix parameter (DIST_CUTOFF = 0.15), OTU clustering parameters (CLUST_ALGO = “average’; OTU_CUTOFF = “0.03”), singletons removal (NSEQ = 1; BYGROUP = false), and distance between sequences (CALC = “onegap”; COUNTENDS = “F”, OUT_TYPE = “square”) was set as shown. This protocol was repeated with the addition of Ion Torrent raw reads from the study published by Vaksmaa and colleagues (2017). In order to normalize for the different sequencing depths between the two data sets, the samples were subjected to a random subsampling at a depth of 10,000 reads. All OTUs with “methylo-” present in their classification were extracted and further identified as the methanotrophic OTUs in the data set.

### Statistical and computational analysis

Subsequent analysis was performed within R version 3.4.1 (R Core Team, 2012). Count data was normalized to relative abundances for all analysis. Analyses were performed with the R package *vegan (*Oksanen et al., 2015). Shannon diversity was calculated using the *diversity* function and Bray-Curtis dissimilarity matrices were generated with *vegdist*. Chao1 estimates were performed using the *chao1* function from the *rareNMtests* package (Cayuela and Gotelli, 2014). Permutational multivariate analyses of variance (PERMANOVA) were performed using the *adonis2* function and classic multidimensional scaling was performed using *cmdscale*. The main drivers of differences in the microbial community composition between paddy field and meadow samples were identified using a random forest classifier from the R package *randomForest (*Liaw and Wiener, 2002).

## Results

### CH_4_ fluxes and Bacterial community structure and composition

The mean CH_4_ flux from the paddy field (0.67 ± 1.76 mg C m^−2^ hr^−2^) and meadow (−0.65 ± 1.59 mg C m^−2^ hr^−2^) did not statistically differ from one another (t = −1.23; p = 0.251; Figure 1). 16S rRNA amplicon sequencing yielded a total of 958,089 reads. On average, each sample contained 12,898 ± 17,506 reads (Table S1). No significant differences were observed between paddy and meadow samples for either species richness (Chao1; t = −0.59; p = 0.574; mean in meadow = 3673 species; mean in paddy = 4260 species) or the sampling depth (t = −0.43, p = 0.67; mean in meadow = 10,179 reads, mean in paddy = 15,342 reads). However, the composition of the total microbial community in paddy and meadow soils significantly differed (PERMAnova; F = 2.27, p = 0.03; Figure 2A). A random forest classifier was used to identify the main bacterial family driving the difference between paddy and meadow soils which was the *Fimbriimonadaceae* that belongs to the phylum Armatimonadetes (formerly OP_10_; Figure 2B). *Fimbriimonadaceae* was significantly more abundant in paddy field soils compared to meadow soils (Figure 2B; t = −4.16; p = 0.001). Furthermore, the bacterial community at the family level was found to be more diverse in the paddy field compared to the meadow (Shannon diversity (H’); t = −3.28; p = 0.005; mean meadow = 3.39; mean paddy field = 3.89; Figure 3).

**Figure 1:**
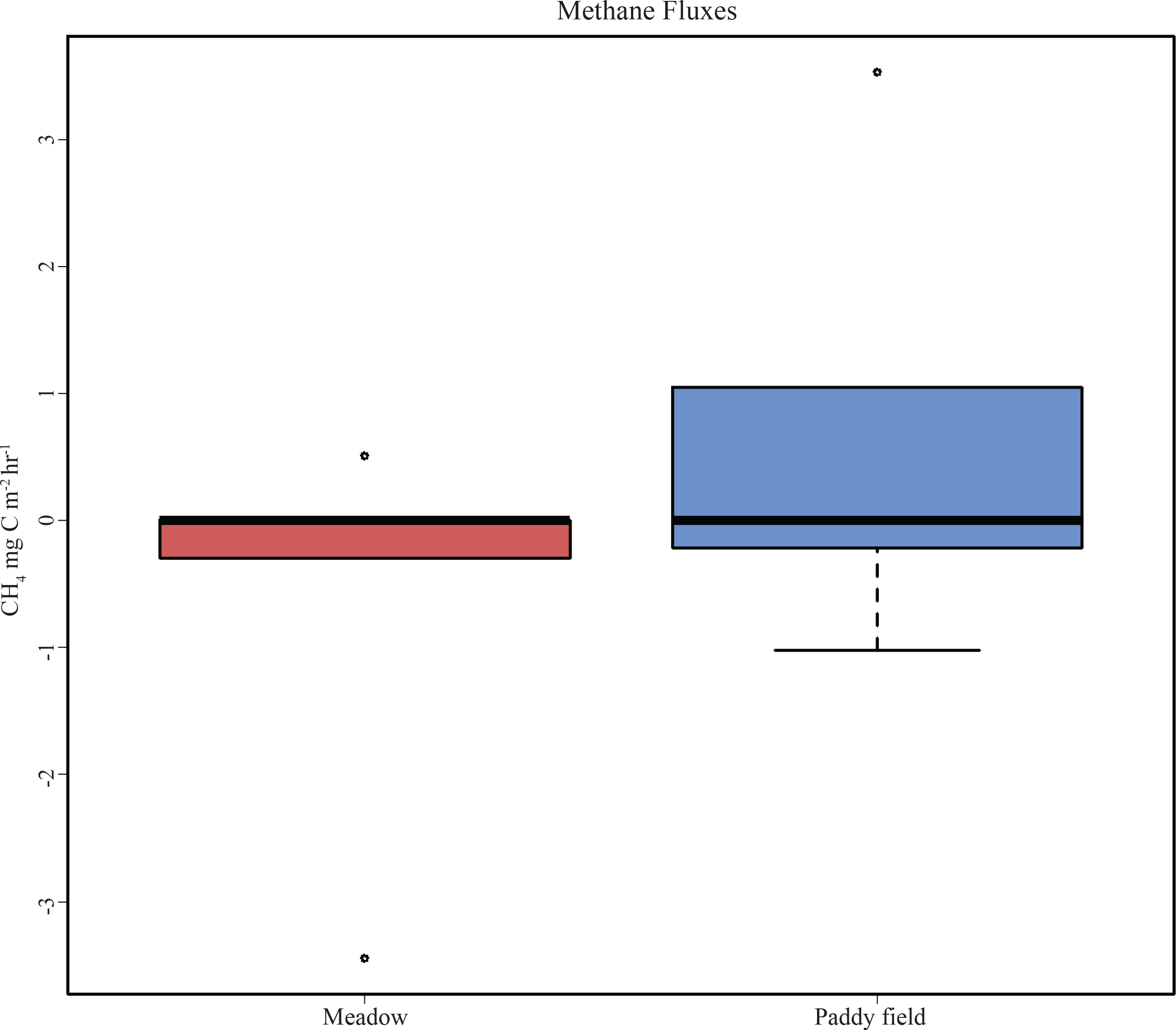
Box-and-whisker plot of methane fluxes in the meadow (**red**) and the paddy field (**blue**) field (t = −1.23; p = 0.251).

**Figure 2:**
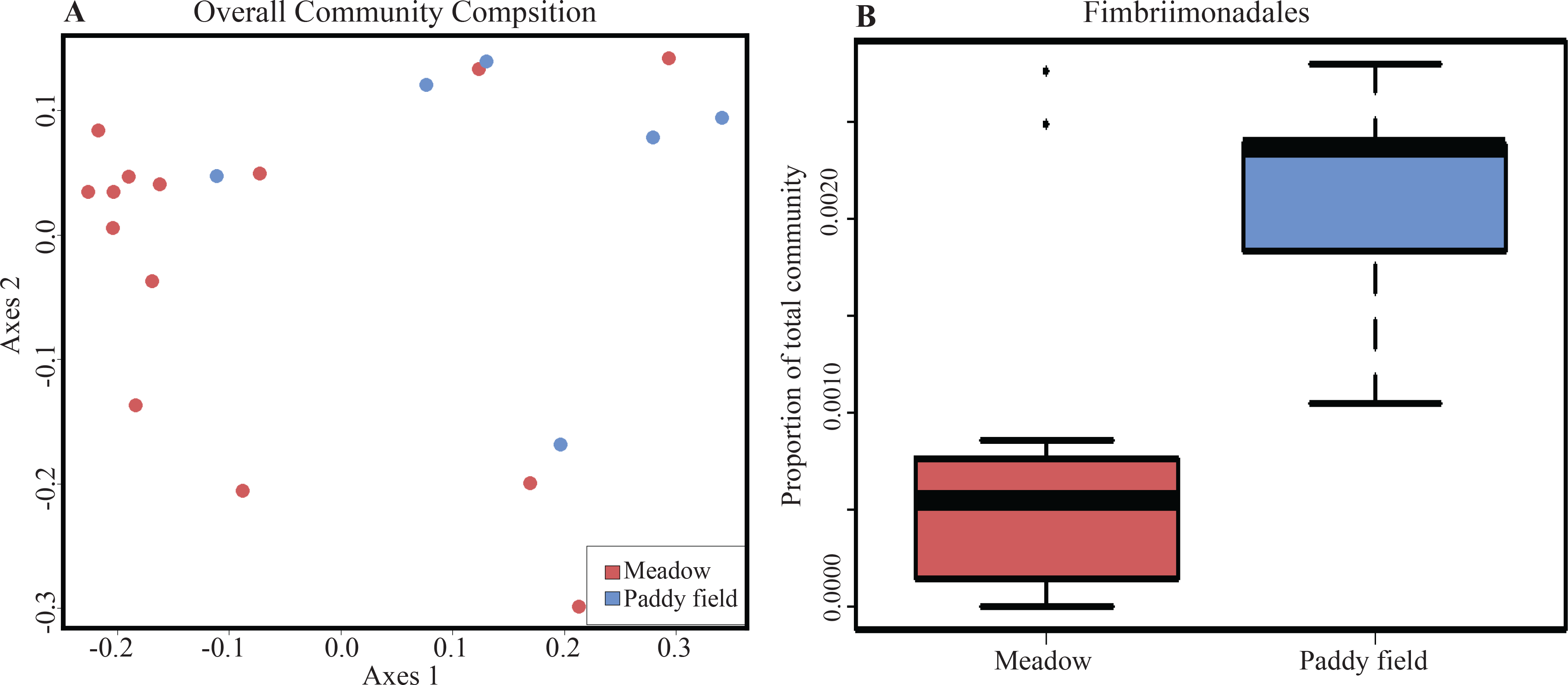
Total bacterial community analysis between the paddy field and the meadow. **A**: MDS plot showing the differences in total bacterial community (PERMAnova, F = 2.27, p = 0.03). Red and blue circles represent the meadow and paddy field samples, respectively. **B**: Relative abundance of *Fimbriimonadales*. Red and blue box- and whisker plots represents meadow and paddy field, respectively.

**Figure 3:**
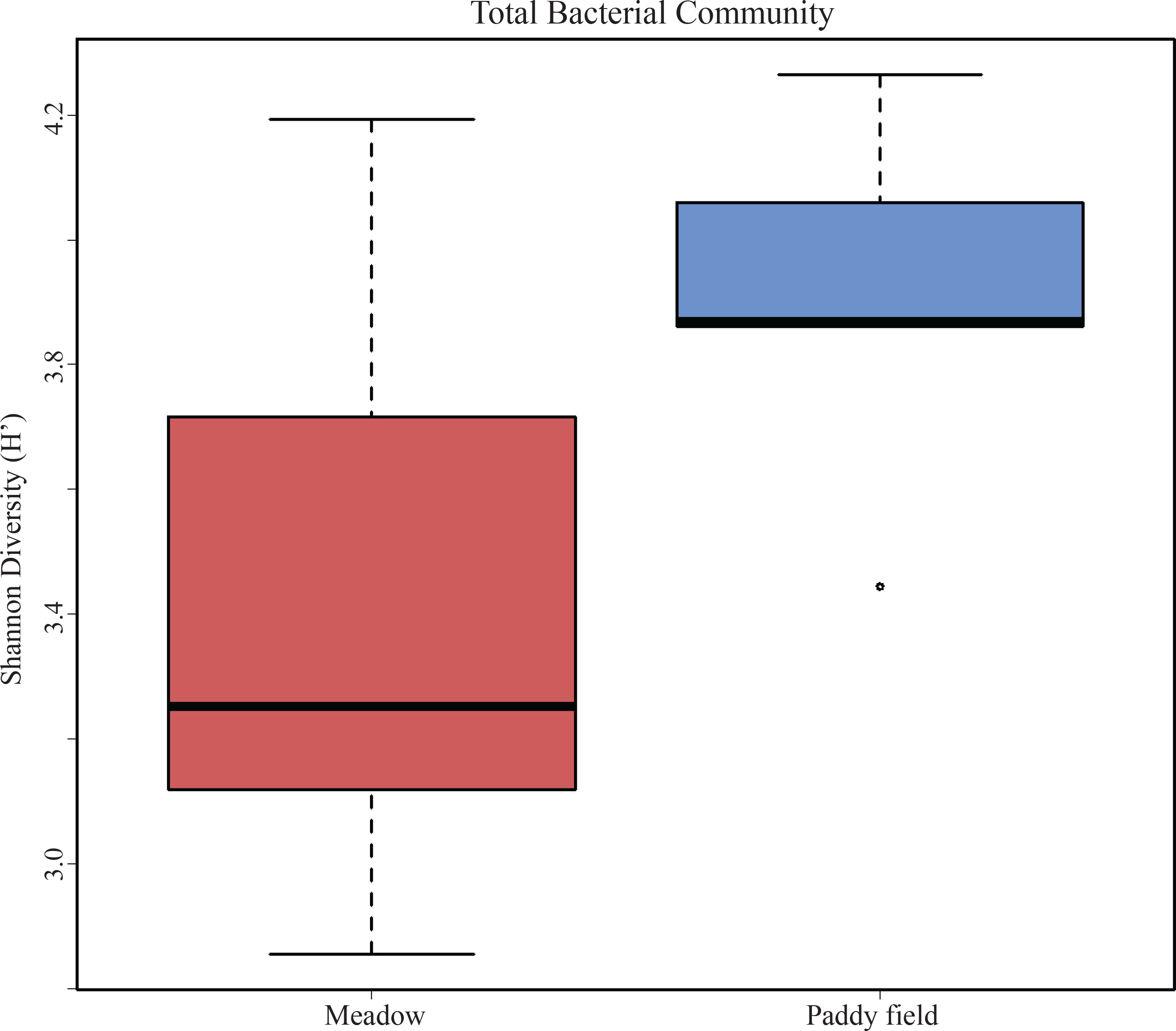
Box-and-whisker plot of total bacterial diversity between paddy field (**blue**) and meadow (**red**) (t-test, t = −3.28, p = 0.005).

### Methylotrophic community

A total of 599 OTUs in the dataset were classified as methanotrophs originating from the *Proteobacteria, Verrucomicrobia* and NC10 phyla. The composition of methanotrophic community differed significantly between the paddy and the meadow soils (PERMAnova; F = 2.63, p = 0.041; Figure 4A). The most abundant methylotroph was affiliated with the family Methyloligellaceae and made up on average 21% of the methylotrophic community. Members of the families Methyloligellaceae and Methylomirabilaceae, were the top two drivers distinguishing the methylotrophic community between meadow and paddy field soils (Figure 4B-C). In general, all methanotroph family abundances were higher in paddy field soils compared to meadow (Figure 5; t = 2.53; p = 0.039).

**Figure 4:**
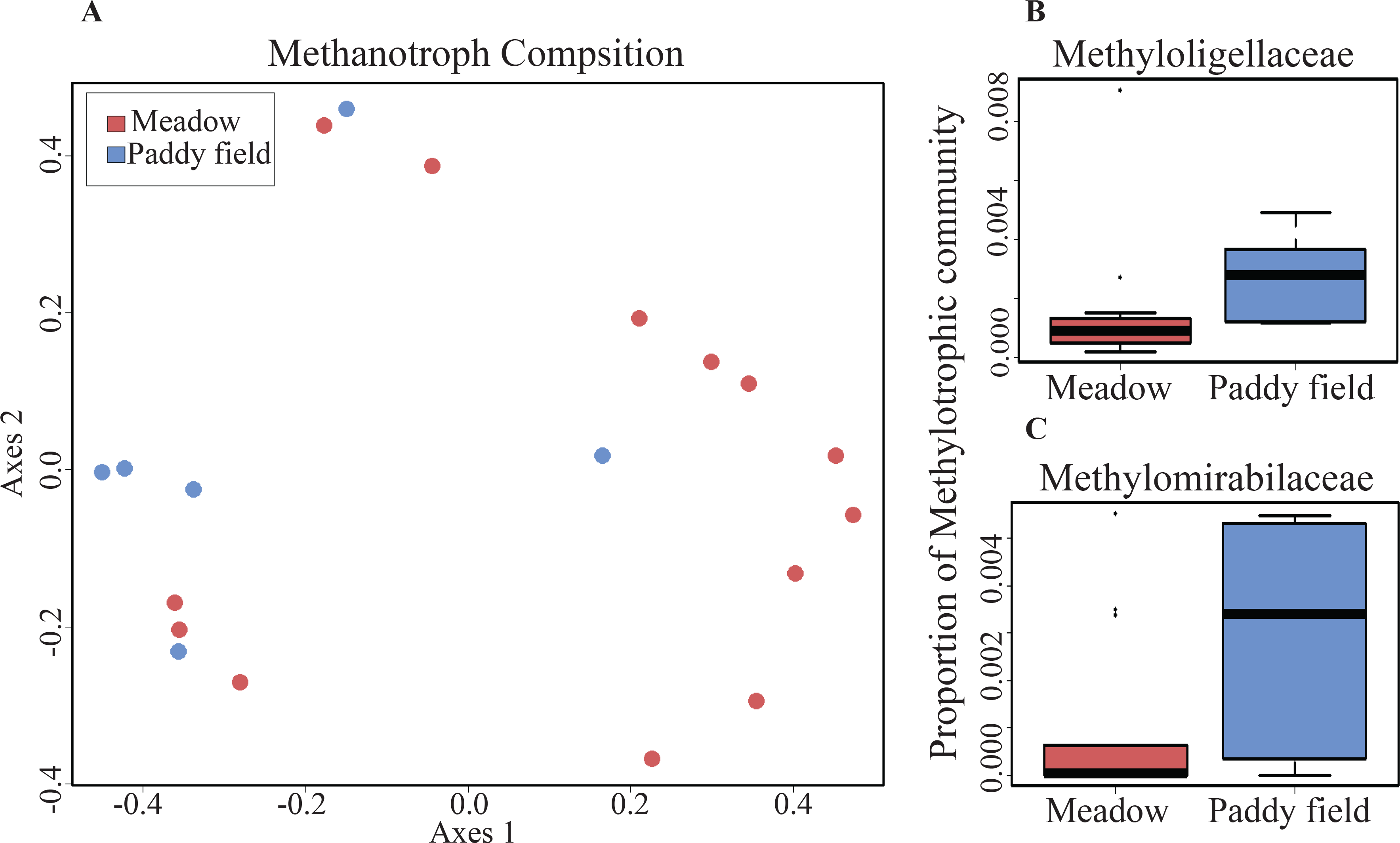
Methanotrophic community analysis between the paddy field and the meadow. **A**: MDS plot of each sampling site showing the difference in the methanotrophic community (PERMAnova, F = 2.63, p = 0.041). Red circles represent the samples within meadow and blue circles represent samples within paddy field. **B and C**: Box-and-whisker plot of top two families that drive the difference in the methylotrophic community between the paddy field (**blue**) than the meadow (**red**).

**Figure 5:**
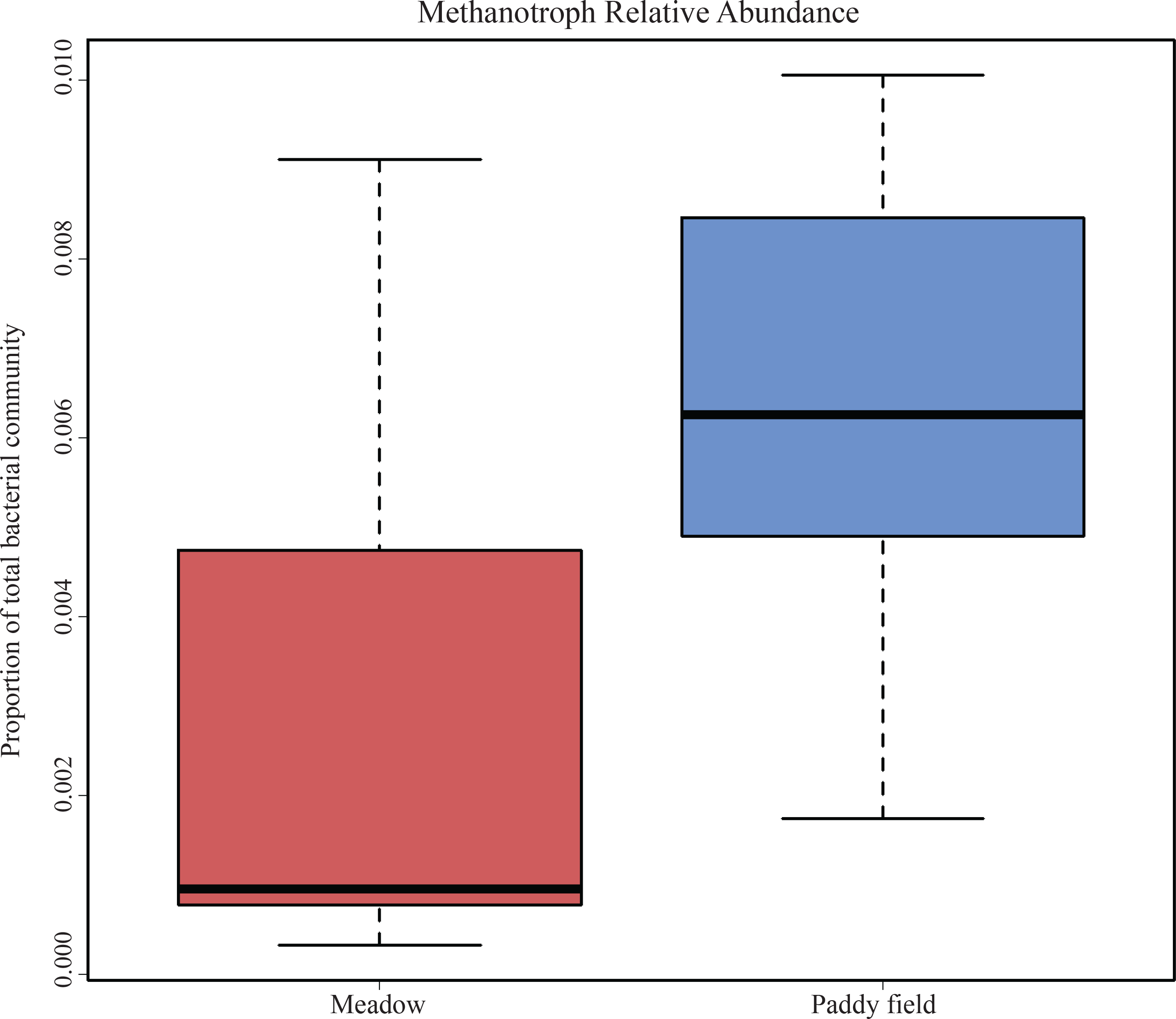
Box-and-whisker plot of difference in relative MOB abundance between the paddy field (**blue**) and the meadow (**red**) (PERMAnova; F = 2.63, p = 0.041).

### Spatial variation of Methanotrophs between paddy fields

In order to examine whether any large-scale spatial variation exists in the methanotrophic community, we compared the data obtained from this study (edge of the paddy field) to the data published by Vaksmaa and colleagues, which were obtained from samples taken from the center of a neighboring paddy field (Vaksmaa et al., 2017). The methanotrophic community in the center of the paddy field more closely resembled the community at the edge of the paddy field than the meadow, however there were differences within the methanotrophic community (Figure 6A; F = 3.91; p = 0.006). Members of *Methylomonaceae* family were significantly more abundant at the center of the neighboring paddy field (Figure 6B;t = 5.24; p = 0.0001). In addition, the *Methylomonaceae* family was found to be the main driver of the differences observed between the two methanotrophic community between the two locations (Figure 6C).

**Figure 6:**
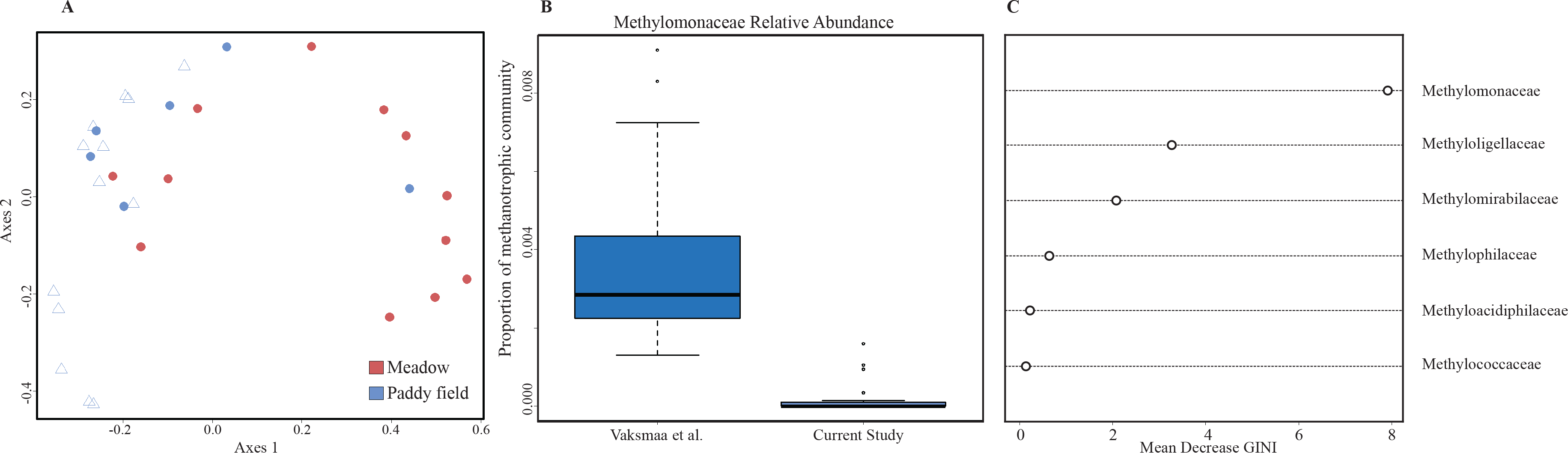
Investigation of spatial variation in methanotrophic community. **A:** MDS plot of each sampling site showing the difference in the methanotrophic community. The meadow (**red**) and paddy field (**blue**) samples of the current study is represented by circles while the paddy field samples from Vaksmaa et al. is represented by blue triangles. **B:** Box-and-whisker plot of relative abundance of Methylomonaceae family between the soil samples at the center of a neighbouring paddy field (Vaksmaa et al., 2017) and the soil from the edge of the paddy field (current study). **C:** Top six families of methanotrophs driving the differences between the two data sets.

## Discussion

The objective of this study was to survey differences in the microbial communities and more specifically the methanotrophic bacteria between a cultivated wetland and an adjacent upland meadow soil, conventionally recognized as strong sources and sinks of CH_4_, respectively. Despite the traditional classification, we observed that the rice paddy field and its neighboring meadow used here are highly variable in their CH_4_ fluxes (Figure 1). This, however, is a single time point and not a representative of possible seasonal fluxes from these fields. It has been reported that during the flooding period of the paddy field, in which no plants are sown, CH_4_ is almost exclusively emitted into the atmosphere through ebullition with fluxes as high as 27.73 mg·g^−2^·h^−1^ (Yuan *et al*., 2018; Holzapfel-Pschorn & Seiler, 1986). After rice cultivation, up to 90% of the observed CH_4_ emissions into the atmosphere are due to diffusion through aerenchyma of the plants. In this study, CH_4_ fluxes and samples were measured from the bulk soil of the paddy field to avoid any possible affect from rice plant or roots on the microbial community or flux data. This could potentially explain the relatively low CH_4_ fluxes observed from the paddy as most of the CH_4_ at the time of sampling would be emitted through the aerenchyma of the rice plants. Additionally, it has been reported that soil amendment by organic fertilization increases the availability of C substrates to methanogens and increases CH_4_ production (Yuan *et al*., 2018; Plaza-Bonilla *et al*., 2014; Mohanty *et al*., 2006). This effect, however, is not observed from the bulk soil in the current study. Furthermore, the paddy field was found to be low in C content (1.4 ± 0.1 %), which would be an indication of low substrate availability for microbial activities. CH_4_ emissions from bulk soil during the maturing stage of rice growth, which is the stage in which the fluxes were measured in this study, are reported to be below 2 mg·g^−2^·h^−1^ (Yuan *et al*., 2018; Vaskmaa et al., 2016).

Despite CH_4_ fluxes being similar between the two sites, we found significant differences in the total bacterial communities’ structure. Since the paddy field is plowed, fertilized and planted homogenously with rice, we hypothesized for the total bacterial composition to be established in a more homogenous manner relative to the meadow. What we found was that the total bacterial community was different from its neighboring meadow with high bacterial heterogeneity observed in both environments (i.e., highly variable; Figure 2A). Despite the variability, we found that the main family driving the difference between the two soils was Fimbriimonadaceae within the Armatimonadetes phylum (Figure 2B). Sequences classified as Armatimonadetes have been obtained by culture-independent methods from various environments including aerobic and anaerobic wastewater treatment processes, hypersaline microbial mats and subsurface geothermal water streams, as well as various rhizospheres (Portillo and Gonzalez 2009; Lee *et al*., 2011; Tamaki *et al*., 2011). The recently isolated *Fimbriimonas ginsengisoli* within the Fimbriimonadia class was described as strictly aerobic, Gram-negative, meso- and neutrophilic strain with the ability to grow on peptone, casamino acids and yeast extract (Im *et al*., 2012). In another study, genera affiliated with the Fimbriimonadaceae grew better in biofilms cultured using a flow incubator with supplied inorganic nitrogen (N) conditions compared to deficient N (Li *et al*., 2017). Since the paddy field more similarly resembles an N condition, it could explain Fimbriimonadaceae’s higher abundance compared to the meadow and moreover, the higher diversity of microbial community observed in the paddy field (Figure 3). However, any role that this family of Armatimonadetes would play in paddy soil N cycling requires further investigation.

Previous research on paddy fields in Vercelli have shown that MOB communities, even in very closely located fields with nearly identical agricultural treatments show significantly different patterns (Ho *et al*., 2011, Lüke *et al*., 2010). This difference in the MOB community was assumed to be a consequence of variance in CH_4_ fluxes (Smith *et al*., 2016; Yuan *et al*., 2014; Holzapfel-Pschorn & Seiler, 1986). However, difference in CH_4_ fluxes could not account for the observations made in this study, particularly regarding the differences observed in the methylotrophic community in the current study (Figure 4A). Furthermore, the most abundant methylotrophs in the paddy field were classified as members of the proposed family “Methyloligellaceae” within the Rhizobiales order (Figure 4B) and Methylomirabilaceae family (Figure 4C).

To date, two separate isolates have been cultivated from Ural saline environments classified as *Methyloligella halotolerans* gen. nov., sp. nov. and *Methyloligella solikamskensis* sp. nov. (Doronina *et al*., 2013). These obligate methylotrophic isolates within the Rhizobiales order are strictly aerobic, Gram negative, non-motile rods that utilize the serine pathway for carbon assimilation. Although environmental sequences belonging to the Methyloligellaceae family have been previously found in agricultural soil (Ceja-Navarro *et al*., 2010), not much is known about the role these alphaproteobacterial methylotrophs play in this environment. Fertilizer application has been shown to have an inhibitory effect on type II methanotrophs in rice fields, while stimulating type I MOB (Mohanty *et al*., 2006). In the current study, we found the Methyloligellaceae family were more abundant in the paddy field compared to the meadow. Without information about their genetics or physiology, it is challenging to infer their possible role in carbon (C) cycling within the paddy field. However, being the most abundant OTU in our data set by an order of magnitude, it is likely that members of this group are key players in this paddy field.

Studies that investigated the presence of methanotrophs in cultivated wetlands have reported the presence of both type I and type II *Methylocaldum*-like and *Methyocystis*-like *pmoA* genes in high abundance (Collet *et al*., 2015; Zheng *et al*, 2008; Asakawa *et al*., 2008). Therefore, we expected the methanotrophic community to be present in higher abundance in the paddy field compared to the meadow (Figure 5). However, we did not find any classical type II methanotrophs (*Methylocystis*- and *Methylosinus*-like) reported to be present in high abundance in cultivated wetlands (Lüke *et al*., 2010; Zheng *et al*., 2008; Shrestha *et al*., 2008). Although multiple factors are in play, an enriched MOB community within the paddy field could be a result of higher indigenous CH_4_ available as substrate when compared to the meadow. This would in turn result in higher rates of CH_4_ oxidation and overall, a higher relative abundance of methanotrophs. In future research, it would be critical to monitor the changes in the MOB community structure during the conversion of a meadow into a paddy field to see what fluctuations occur in the MOB community after a drastic change in the cultivation regiment.

Lastly, due to the highly variable CH_4_ fluxes throughout the paddy field, we wanted to investigate if sampling location influenced the abundance of methanotrophs. Previous studies on the spatial variation of methane fluxes within paddy fields have demonstrated that fluxes can vary significantly even within the same field, thus making extrapolation to larger areas from point samples challenging (Oo *et al*., 2015; Sass *et al*., 2002; Krause *et al*., 2009; Spokas *et al*., 2013). In order to compare the community composition of methanotrophs between two neighboring paddy fields, we incorporated the data published by Vaksmaa and colleagues into our analyses (Vaskmaa *et al*., 2017; Figure 6A-C). We found that while the methanotrophic community from the Vaskmaa et al. study resembled that of our paddy soil samples, there was a significant difference in relative abundance of certain methanotrophic families between the two sampling locations (Figure 6A-B). More specifically, members of the *Methylomonaceae* family were found in much higher relative abundance in the previous study (Figure 6B) and were found to be the main family, driving the differences observed between the two data sets (Figure 6C). Unfortunately, there is no CH_4_ flux data available to compare between the two study sites and whether more indigenous CH_4_ availability is the reason behind this difference. Regardless, this finding suggests that the methanotrophic community may be heterogeneously distributed across neighboring paddy fields, possibly resulting in differences in CH_4_ oxidation capabilities. As indicated by others, designing a strong experiment which pairs soil samples for microbial community analysis, in situ measurements of important environmental factors (i.e., pore water pH, inorganic compound concentrations, etc) with CH_4_ flux data may reveal drivers of the heterogeneity of the paddy field methanotrophic community (Hester et al., 2018).

## Conclusion

Typically, paddy fields are regarded as CH_4_ sources. These CH_4_ emissions stem from a combination of higher fertilizer inputs, which result in increased organic matter deposited by the rice plant into the rhizosphere, providing substrate for methanogenesis (i.e., higher methanogenic activity). In the current study, we find that the paddy field on average was a source (positive CH_4_ fluxes) and the meadow a sink (negative CH_4_ fluxes), though there was no statistically significant difference between the two locations due to the high variability of the flux measurements. Based on the data collected and previous studies, we propose two possible working hypotheses responsible for the observations made by this study (Figure 7). The observed low levels of CH_4_ emissions in the paddy field could be due to high turnover rates of CH_4_. This is indicated by the high relative abundance of the MOB community comprised mainly of type II methanotrophs affiliated with the *Methyloligellaceae* family, and complies with previous studies (Barbosa *et al*., 2018; Yuan *et al*., 2018). Alternatively, the low flux could stem from a lower initial CH_4_ production within the soil due to decreased various root exudates, a lower redox potential, or the provision of methanogenic substrates by heterotrophic bacteria (Mayer & Conrad, 1990; Aulakh *et al*., 2001). Consequently, due to the high variability within and between wetland and upland soils, caution should be exercised when making extrapolated predictions of CH_4_ emissions.

**Figure 7:**
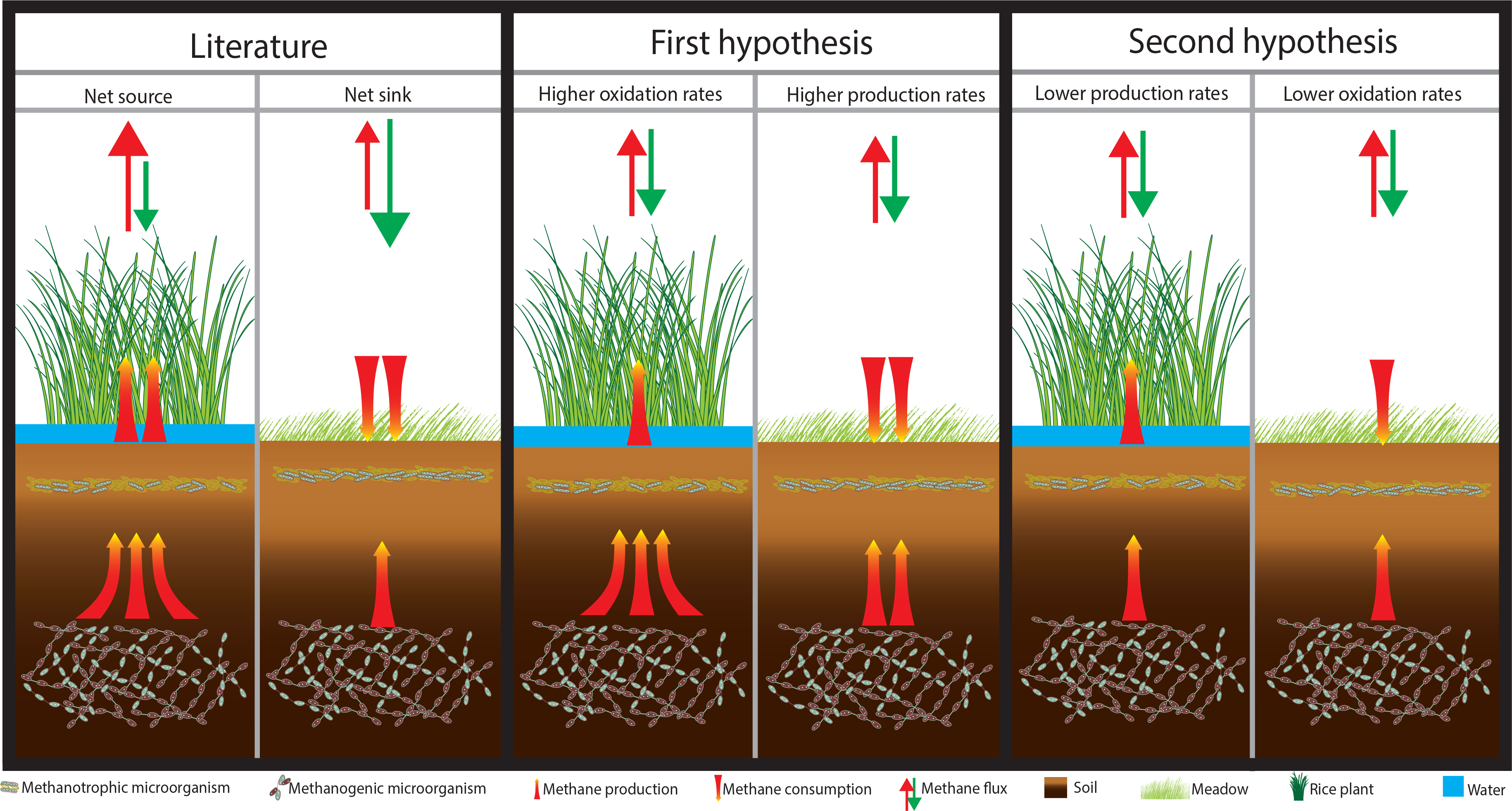
Hypotheses based on the data collected and previous. **A:** The paddy field and the meadow act as a CH_4_ source and a sink, respectively. **B and C**: Two working models based on the findings of the current study on the paddy field and meadow not fulfilling their role as a CH_4_ source and a sink.

## Acknowledgement

Research was supported by ERC ecomom 339880 and siam OCW/NWO 024002002. We thank the CREA-Rice Research Unit of Vercelli (Italy) for assistance during sampling.

## References

Asakawa, S., & Kimura, M. (2008). Comparison of bacterial community structures at main habitats in paddy field ecosystem based on DGGE analysis. Soil Biology and Biochemistry, 40(6), 1322– 1329.

Aselmann, I., & Crutzen, P. (1989). Global distribution of natural freshwater wetlands and rice paddies, their net primary productivity, seasonality and possible methane emissions. Journal of Atmospheric chemistry, 8(4), 307–358.

Aulakh, M. S., Wassmann, R., Bueno, C., & Rennenberg, H. (2001). Impact of root exudates of different cultivars and plant development stages of rice (Oryza sativa L.) on methane production in a paddy soil. Plant and Soil, 230(1), 77–86.

Barbosa, P. M., Farjalla, V. F., Melack, J. M., Amaral, J. H. F., da Silva, J. S., & Forsberg, B. R. (2018). High rates of methane oxidation in an Amazon floodplain lake. Biogeochemistry, 137(3), 351–365.

Bodelier, P. (2011). Toward understanding, managing, and protecting microbial ecosystems. Frontiers in microbiology, 2, 80.

Cao, M., Gregson, K., & Marshall, S. (1998). Global methane emission from wetlands and its sensitivity to climate change. Atmospheric environment, 32(19), 3293–3299.

Ceja-Navarro, J. A., Rivera-Orduna, F. N., Patino-Zúniga, L., Vila-Sanjurjo, A., Crossa, J., Govaerts, B., & Dendooven, L. (2010). Phylogenetic and multivariate analyses to determine the effects of different tillage and residue management practices on soil bacterial communities. Applied and Environmental Microbiology, 76(11), 3685–3691.

Collet, S., Reim, A., Ho, A., & Frenzel, P. (2015). Recovery of paddy soil methanotrophs from long term drought. Soil Biology and Biochemistry, 88, 69–72.

Conrad, R. (2009). The global methane cycle: recent advances in understanding the microbial processes involved. Environmental Microbiology Reports, 1(5), 285–292.

Danilova, O. V., Suzina, N. E., Van De Kamp, J., Svenning, M. M., Bodrossy, L., & Dedysh, S. N. (2016). A new cell morphotype among methane oxidizers: a spiral-shaped obligately microaerophilic methanotroph from northern low-oxygen environments. The ISME journal, 10(11), 2734.

Dean, J. F., Middelburg, J. J., Röckmann, T., Aerts, R., Blauw, L. G., Egger, M.,… Rasigraf, O. (2018). Methane feedbacks to the global climate system in a warmer world. Reviews of Geophysics.

Dedysh, S. N., Berestovskaya, Y. Y., Vasylieva, L. V., Belova, S. E., Khmelenina, V. N., Suzina, N. E.,… Zavarzin, G. A. (2004). *Methylocella tundrae* sp. nov., a novel methanotrophic bacterium from acidic tundra peatlands. International Journal of Systematic and Evolutionary Microbiology, 54(1), 151–156.

DeSantis, T. Z., Brodie, E. L., Moberg, J. P., Zubieta, I. X., Piceno, Y. M., & Andersen, G. L. (2007). High-density universal 16S rRNA microarray analysis reveals broader diversity than typical clone library when sampling the environment. Microbial ecology, 53(3), 371–383.

Dlugokencky, E., Masaire, K., Lang, P., Tans, P., Steele, L., & Nisbet, E. (1994). A dramatic decrease in the growth rate of atmospheric methane in the northern hemisphere during 1992. Geophysical Research Letters, 21(1), 45–48.

Dobbie, K., Smith, K., Prieme, A., Christensen, S., Degorska, A., & Orlanski, P. (1996). Effect of land use on the rate of methane uptake by surface soils in northern Europe. Atmospheric environment, 30(7), 1005–1011.

Doronina, N. V., Poroshina, M. N., Kaparullina, E. N., Ezhov, V. A., & Trotsenko, Y. A. (2013). *Methyloligella halotolerans* gen. nov., sp. nov. and *Methyloligella solikamskensis* sp. nov., two non-pigmented halotolerant obligately methylotrophic bacteria isolated from the Ural saline environments. Systematic and applied microbiology, 36(3), 148–154.

Ettwig, K. F., Butler, M. K., Le Paslier, D., Pelletier, E., Mangenot, S., Kuypers, M. M.,… De Beer, D. (2010). Nitrite-driven anaerobic methane oxidation by oxygenic bacteria. Nature, 464(7288), 543.

Fjellbirkeland, A., Torsvik, V., & Øvreås, L. (2001). Methanotrophic diversity in an agricultural soil as evaluated by denaturing gradient gel electrophoresis profiles of *pmoA, mxaF* and 16S rDNA sequences. Antonie van Leeuwenhoek, 79(2), 209–217.

Flato, G., Marotzke, J., Abiodun, B., Braconnot, P., Chou, S. C., Collins, W. J.,… Eyring, V. (2013). Evaluation of climate models. In: climate change 2013: the physical science basis. Contribution of working group I to the fifth assessment report of the intergovernmental panel on climate change. Climate Change 2013, 5, 741–866.

Garcia, J.-L., Patel, B. K., & Ollivier, B. (2000). Taxonomic, phylogenetic, and ecological diversity of methanogenic Archaea. Anaerobe, 6(4), 205–226.

Haroon, M. F., Hu, S., Shi, Y., Imelfort, M., Keller, J., Hugenholtz, P.,… Tyson, G. W. (2013). Anaerobic oxidation of methane coupled to nitrate reduction in a novel archaeal lineage. Nature, 500(7464), 567.

Henckel, T., Roslev, P., & Conrad, R. (2000). Effects of O2 and CH4 on presence and activity of the indigenous methanotrophic community in rice field soil. Environmental Microbiology, 2(6), 666–679.

Hester, E. R., Harpenslager, S. F., van Diggelen, J. M., Lamers, L. L., Jetten, M. S., Lüke, C.,… & Welte, C. U. (2018). Linking nitrogen load to the structure and function of wetland soil and rhizosphere microbial communities. Msystems, 3(1), e00214–17.

Ho, A., Lüke, C., & Frenzel, P. (2011). Recovery of methanotrophs from disturbance: population dynamics, evenness and functioning. The ISME journal, 5(4), 750.

Holzapfel-Pschorn, A., & Seiler, W. (1986). Methane emission during a cultivation period from an Italian rice paddy. Journal of Geophysical Research: Atmospheres, 91(D11), 11803–11814.

Horz, H.-P., Yimga, M. T., & Liesack, W. (2001). Detection of methanotroph diversity on roots of submerged rice plants by molecular retrieval of *pmoA, mmoX, mxaF*, and 16S rRNA and ribosomal DNA, including *pmoA*-based terminal restriction fragment length polymorphism profiling. Applied and Environmental Microbiology, 67(9), 4177–4185.

Hutchinson, G., & Livingston, G. (2001). Vents and seals in non-steady-state chambers used for measuring gas exchange between soil and the atmosphere. European Journal of Soil Science, 52(4), 675–682.

Im, W.-T., Hu, Z.-Y., Kim, K.-H., Rhee, S.-K., Meng, H., Lee, S.-T., & Quan, Z.-X. (2012). Description of *Fimbriimonas ginsengisoli* gen. nov., sp. nov. within the Fimbriimonadia class nov., of the phylum Armatimonadetes. Antonie van Leeuwenhoek, 102(2), 307–317.

Janssen, P. H. (2006). Identifying the dominant soil bacterial taxa in libraries of 16S rRNA and 16S rRNA genes. Applied and Environmental Microbiology, 72(3), 1719–1728.

Johnston, H. S. (1984). Human effects on the global atmosphere. Annual Review of Physical Chemistry, 35(1), 481–505.

Kalyuzhnaya, M., Yang, S., Rozova, O., Smalley, N., Clubb, J., Lamb, A.,… Bringel, F. (2013). Highly efficient methane biocatalysis revealed in a methanotrophic bacterium. Nature communications, 4, 2785.

Kits, K. D., Klotz, M. G., & Stein, L. Y. (2015). Methane oxidation coupled to nitrate reduction under hypoxia by the Gammaproteobacterium *Methylomonas denitrificans*, sp. nov. type strain FJG1. Environmental Microbiology, 17(9), 3219–3232.

Klindworth, A., Pruesse, E., Schweer, T., Peplies, J., Quast, C., Horn, M., & Glöckner, F. O. (2013). Evaluation of general 16S ribosomal RNA gene PCR primers for classical and next-generation sequencing-based diversity studies. Nucleic acids research, 41(1), e1–e1.

Li, S., Peng, C., Wang, C., Zheng, J., Hu, Y., & Li, D. (2017). Microbial succession and nitrogen cycling in cultured biofilms as affected by the inorganic nitrogen availability. Microbial ecology, 73(1), 1–15.

Lin, X., Wang, S., Hu, Y., Luo, C., Zhang, Z., Niu, H., & Xie, Z. (2015). Experimental warming increases seasonal methane uptake in an alpine meadow on the Tibetan Plateau. Ecosystems, 18(2), 274–286.

Livingston, G. P., Hutchinson, G. L., & Spartalian, K. (2005). Diffusion theory improves chamber-based measurements of trace gas emissions. Geophysical Research Letters, 32(24).

Lüke, C., Krause, S., Cavigiolo, S., Greppi, D., Lupotto, E., & Frenzel, P. (2010). Biogeography of wetland rice methanotrophs. Environmental Microbiology, 12(4), 862–872.

Mayer, H. P., & Conrad, R. (1990). Factors influencing the population of methanogenic bacteria and the initiation of methane production upon flooding of paddy soil. FEMS Microbiology Ecology, 6(2), 103–111.

Miyata, A., Leuning, R., Denmead, O. T., Kim, J., & Harazono, Y. (2000). Carbon dioxide and methane fluxes from an intermittently flooded paddy field. Agricultural and Forest Meteorology, 102(4), 287–303.

Mohanty, S. R., Bodelier, P. L., Floris, V., & Conrad, R. (2006). Differential effects of nitrogenous fertilizers on methane-consuming microbes in rice field and forest soils. Applied and Environmental Microbiology, 72(2), 1346–1354.

Mosier, A., Duxbury, J., Freney, J., Heinemeyer, O., Minami, K., & Johnson, D. (1998). Mitigating agricultural emissions of methane. Climatic Change, 40(1), 39–80.

Mosier, A., Schimel, D., Valentine, D., Bronson, K., & Parton, W. (1991). Methane and nitrous oxide fluxes in native, fertilized and cultivated grasslands. Nature, 350(6316), 330.

Oo, A. Z., Win, K. T., & Bellingrath-Kimura, S. D. (2015). Within field spatial variation in methane emissions from lowland rice in Myanmar. SpringerPlus, 4(1), 145.

Op den Camp, H. J., Islam, T., Stott, M. B., Harhangi, H. R., Hynes, A., Schouten, S.,… Dunfield, P. F. (2009). Environmental, genomic and taxonomic perspectives on methanotrophic Verrucomicrobia. Environmental Microbiology Reports, 1(5), 293–306.

Padilla, C. C., Bertagnolli, A. D., Bristow, L. A., Sarode, N., Glass, J. B., Thamdrup, B., & Stewart, F. J. (2017). Metagenomic binning recovers a transcriptionally active Gammaproteobacterium linking methanotrophy to partial denitrification in an anoxic oxygen minimum zone. Frontiers in Marine Science, 4, 23.

Paudel, R., Mahowald, N. M., Hess, P. G., Meng, L., & Riley, W. J. (2016). Attribution of changes in global wetland methane emissions from pre-industrial to present using CLM4. 5-BGC. Environmental Research Letters, 11(3), 034020.

Pearman, G., Etheridge, D., De Silva, F., & Fraser, P. (1986). Evidence of changing concentrations of atmospheric CO_2_, N_2_O and CH_4_ from air bubbles in Antarctic ice. Nature, 320(6059), 248.

Pedersen, A. R., Petersen, S. O., & Schelde, K. (2010). A comprehensive approach to soil-atmosphere trace-gas flux estimation with static chambers. European Journal of Soil Science, 61(6), 888– 902.

Plaza-Bonilla, D., Cantero-Martínez, C., Bareche, J., Arrúe, J. L., & Álvaro-Fuentes, J. (2014). Soil carbon dioxide and methane fluxes as affected by tillage and N fertilization in dryland conditions. Plant and Soil, 381(1-2), 111-130.

Pumpanen, J., Kolari, P., Ilvesniemi, H., Minkkinen, K., Vesala, T., Niinistö, S.,… Pihlatie, M. (2004). Comparison of different chamber techniques for measuring soil CO_2_ efflux. Agricultural and Forest Meteorology, 123(3-4), 159-176.

Reeburgh, W. (1993). The role of methylotrophy in the global methane budget. Microbial growth on C-1 compounds, 1–14.

Salipante, S. J., Kawashima, T., Rosenthal, C., Hoogestraat, D. R., Cummings, L. A., Sengupta, D. J.,… Hoffman, N. G. (2014). Performance comparison of Illumina and ion torrent next-generation sequencing platforms for 16S rRNA-based bacterial community profiling. Applied and Environmental Microbiology, 80(24), 7583–7591.

Sanschagrin, S., & Yergeau, E. (2014). Next-generation sequencing of 16S ribosomal RNA gene amplicons. Journal of visualized experiments: JoVE(90).

Sass, R. L., Fisher, F. M., & Andrews, J. A. (2002). Spatial variability in methane emissions from a Texas rice field with some general implications. Global biogeochemical cycles, 16(1).

Schimel, D., Alves, D., Enting, I., Heimann, M., Joos, F., Raynaud, D.,… Ehhalt, D. (1996). Radiative forcing of climate change. Climate change 1995: The science of climate change, 65– 131.

Shen, L.-d., Liu, S., Huang, Q., Lian, X., He, Z.-f., Geng, S.,… Xu, X.-y. (2014). Evidence for the cooccurrence of nitrite-dependent anaerobic ammonium and methane oxidation processes in a flooded paddy field. Applied and Environmental Microbiology, 80(24), 7611–7619.

Shrestha, M., Abraham, W. R., Shrestha, P. M., Noll, M., & Conrad, R. (2008). Activity and composition of methanotrophic bacterial communities in planted rice soil studied by flux measurements, analyses of *pmoA* gene and stable isotope probing of phospholipid fatty acids. Environmental Microbiology, 10(2), 400–412.

Smith, K., Ball, T., Conen, F., Dobbie, K., Massheder, J., & Rey, A. (2003). Exchange of greenhouse gases between soil and atmosphere: interactions of soil physical factors and biological processes. European Journal of Soil Science, 54(4), 779–791.

Spokas, K., Graff, C., Morcet, M., & Aran, C. (2003). Implications of the spatial variability of landfill emission rates on geospatial analyses. Waste Management, 23(7), 599–607.

Team, R. C. (2013). R: A language and environment for statistical computing.

Tremblay, J., Singh, K., Fern, A., Kirton, E. S., He, S., Woyke, T.,… Tringe, S. G. (2015). Primer and platform effects on 16S rRNA tag sequencing. Frontiers in microbiology, 6, 771.

Vaksmaa, A., Lüke, C., Van Alen, T., Valè, G., Lupotto, E., Jetten, M., & Ettwig, K. (2016). Distribution and activity of the anaerobic methanotrophic community in a nitrogen-fertilized Italian paddy soil. FEMS Microbiology Ecology, 92(12).

Vaksmaa, A., van Alen, T. A., Ettwig, K. F., Lupotto, E., Valè, G., Jetten, M. S., & Lüke, C. (2017). Stratification of diversity and activity of methanogenic and methanotrophic microorganisms in a nitrogen-fertilized Italian paddy soil. Frontiers in microbiology, 8, 2127.

Wahlen, M. (1993). The global methane cycle. Annual Review of Earth and Planetary Sciences, 21(1), 407–426.

Welte, C. U., Rasigraf, O., Vaksmaa, A., Versantvoort, W., Arshad, A., Op den Camp, H. J.,… Reimann, J. (2016). Nitrate-and nitrite-dependent anaerobic oxidation of methane. Environmental Microbiology Reports, 8(6), 941–955.

Whittenbury, R., Phillips, K., & Wilkinson, J. (1970). Enrichment, isolation and some properties of methane-utilizing bacteria. Microbiology, 61(2), 205–218.

Wu, M. L., Ettwig, K. F., Jetten, M. S., Strous, M., Keltjens, J. T., & van Niftrik, L. (2011). A new intra-aerobic metabolism in the nitrite-dependent anaerobic methane-oxidizing bacterium *Candidatus* ‘Methylomirabilis oxyfera’: Portland Press Limited.

Xiao, X., Boles, S., Liu, J., Zhuang, D., Frolking, S., Li, C.,… Moore III, B. (2005). Mapping paddy rice agriculture in southern China using multi-temporal MODIS images. Remote sensing of environment, 95(4), 480–492.

Yan, X., Yagi, K., Akiyama, H., & Akimoto, H. (2005). Statistical analysis of the major variables controlling methane emission from rice fields. Global Change Biology, 11(7), 1131–1141.

Yarza, P., Yilmaz, P., Pruesse, E., Glöckner, F. O., Ludwig, W., Schleifer, K.-H.,… Rosselló-Móra, R. (2014). Uniting the classification of cultured and uncultured bacteria and archaea using 16S rRNA gene sequences. Nature Reviews Microbiology, 12(9), 635.

Yuan, J., Yuan, Y., Zhu, Y., & Cao, L. (2018). Effects of different fertilizers on methane emissions and methanogenic community structures in paddy rhizosphere soil. Science of The Total Environment, 627, 770–781.

Yuan, Q., Pump, J., & Conrad, R. (2014). Straw application in paddy soil enhances methane production also from other carbon sources. Biogeosciences, 11(2), 237–246.

Zheng, Y., Zhang, L.-M., Zheng, Y.-M., Di, H., & He, J.-Z. (2008). Abundance and community composition of methanotrophs in a Chinese paddy soil under long-term fertilization practices. Journal of Soils and Sediments, 8(6), 406–414.

