## Supplementary Table 1, 2 and 3 and supplementary figure 1 for "Comparison of the bacterial and methanotrophic diversities between an Italian paddy field and its neighboring meadow"

**Supplementary figures and tables:**

**Table S1:** Number of sequences obtained from each sample at pre- and post-analysis.

| **Sample** | **Location** | **Pre-analysis** | **Post-analysis** |
| --- | --- | --- | --- |
| 1 | Meadow | 50470 | 22581 |
| 2 | Meadow | 47218 | 9273 |
| 3 | Meadow | 75573 | 30682 |
| 4 | Meadow | 27496 | 5328 |
| 5 | Meadow | 78818 | 33284 |
| 6 | Meadow | 87835 | 19081 |
| 7 | Paddy | 198593 | 77899 |
| 8 | Paddy | 7842 | 1372 |
| 9 | Paddy | 9433 | 1702 |
| 10 | Meadow | 28515 | 5130 |
| 11 | Meadow | 29000 | 5396 |
| 12 | Meadow | 2035 | 366 |
| 13 | Paddy | 37795 | 7099 |
| 14 | Paddy | 40746 | 6909 |
| 15 | Paddy | 30811 | 5974 |
| 16 | Meadow | 14987 | 2788 |
| 17 | Meadow | 48306 | 8884 |
| 18 | Meadow | 18682 | 3755 |
| 19 | Meadow | 21624 | 4440 |
| 20 | Meadow | 28676 | 6036 |
| Total number of sequences | | **958089** | **281043** |

**Table S2:** A brief metadata of studies reporting on wetlands as a CH_4_ source and uplands as a CH_4_ sink environment with contradicting findings.

| References | Environment | Source vs. Sink | Applied Method |
| --- | --- | --- | --- |
| Wetlands |  |  |  |
| Feng *et al.*, 2012 | Paddy field | **Source** | PCR-DGGE and qPCR |
| Banger *et al.*, 2012 | Paddy field | **Source** | 33 published papers on the subject |
| Ma *et al.*, 2010 | Paddy field | **Source** | RFLP of 16S rRNA, pmoA |
| Lee *et al.*, 2014 | Paddy field | **Source** | *pmoA* and *mcrA* transcripts |
| Shrestha *et al.*, 2010 | Paddy field | **Source** | T-RFLP of *pmoA* |
| Ma et al., 2010 | Paddy field | **Source** | qPCR and T-RFLP |
| Hoffmann *et al.*, 2002 | Paddy field | **Source** | T-RFLP of *pmoA* and DGGE |
| Conrad *et al.*, 2007 | Paddy field | **Source** | T-RFLP of 16S rRNA and *mcrA* |
| Bodelier *et al.*, 2000 | Paddy field | **Source** | Radioactive fingerprinting |
| Noll *et al.*, 2008 | Paddy field | **Source** | T-RFLP |
| Eller & Frenzel, 2001 | Paddy field | **Source** | DGGE and FISH |
| Uplands |  |  |  |
| Tate *et al.*, 2007 | Shrubland | **Sink** | Flux measurements |
| Henckel *et al.*, 2000 | Forest | **Sink** | Methane profile and DGGE |
| Knief & Dunfield, 2005 | Upland soil | **Sink** | Molecular techniques and CH_4_ flux |
| Benstead & King, 2001 | Forest | **Sink** | Methane flux analysis |
| Groffman *&* Pouyat., 2009 | Forest & Lawns | **Sink** | Methane flux analysis |
| Kolb *et al.*, 2005 | Forest | **Sink** | Molecular techniques |
| Luo *et al.*, 2013 | Forest | **Sink** | Methane flux analysis |
| Fang *et al.*, 2014 | Meadow | **Sink** | Methane flux analysis |
| Jang *et al.*, 2011 | Forest | **Sink** | Methane flux analysis and T-RFLP |
| Aronson & Helliker, 2010 | Upland soil | **Sink** | Meta-analysis |
| Blankinship *et al.*, 2010 | Grassland | **Sink** | Multifactor analysis |

B

F

E

D

C

A

P	P	P	M1	M2	M3

Paddy	Meadow

**Figure S1:** Sampling sites within paddy field and meadow represented by white circles. The CM1 sampling point was discarded due to poor sequence quality.
